# Human pluripotent stem cell-derived hepatocyte-like cells for hepatitis D virus studies

**DOI:** 10.1101/2023.10.13.561984

**Authors:** Huanting Chi, Bingqian Qu, Angga Prawira, Lars Maurer, Jungen Hu, Rebecca M. Fu, Florian A. Lempp, Zhenfeng Zhang, Dirk Grimm, Xianfang Wu, Stephan Urban, Viet Loan Dao Thi

## Abstract

Current culture systems available for studying hepatitis D virus (HDV) are suboptimal. In this study, we demonstrate that hepatocyte-like cells (HLCs) derived from human pluripotent stem cells (hPSCs) are fully permissive to HDV infection across various tested genotypes. When co-infected with the helper hepatitis B virus (HBV) or transduced to express the HBV envelope protein HBsAg, HLCs effectively secrete infectious progeny virions. We also show that HLCs expressing HBsAg support extracellular spread of HDV, thus providing a valuable platform for testing available anti-HDV regimens. By challenging the cells along the differentiation with HDV infection, we have identified CD63 as a potential HDV/HBV co-entry factor, which was rate-limiting HDV infection in immature hepatocytes. Given their renewable source and the potential to derive hPSCs from individual patients, we propose HLCs as a promising model for investigating HDV biology. Our findings offer new insights into HDV infection and expand the repertoire of research tools available for the development of therapeutic interventions.

**Teaser:** A human stem cell-derived hepatocyte culture model for hepatitis D virus studies

## Introduction

Approximately 5% of chronic hepatitis B virus (HBV) carriers are co- or super-infected with hepatitis D virus (HDV), resulting in an estimated 12 million chronic hepatitis D (CHD) patients worldwide^1^. However, a recent study suggests that these numbers probably represent an underestimation of the true global burden of disease^2^. HBV/HDV co-infection can cause the most aggressive form of viral hepatitis, leading to an accelerated progression of liver dysfunction and disease^3^.

HDV is a circular, single-stranded, negative-sense RNA virus that belongs to the *Kolmioviridae* family. Based on their genome diversity, eight different HDV genotypes (GTs) have been described, each showing distinct geographical distributions^4^. HDV-1 is the most prevalent genotype and endemic worldwide, while infections with genotypes HDV-2 through -8 occur regionally. HDV-2 infections are common in Southeast Asia, while the most diverged genotype HDV-3 is exclusively prevalent in South America. HDV-4 is found in Japan and Taiwan and HDV- 5 through -8 infections are restricted to Africa^5^. Infection with the different GTs can lead to various disease outcomes in the clinic: Compared to HDV-2, both HDV-1 and HDV-3 infections can lead to severe hepatitis with more adverse patient outcomes^6^. Patients infected with HDV-4 usually experience a mild course of liver disease. Patients with an HDV-5 infection appear to have a preferable prognosis and respond better to pegylated interferon alpha (peg-IFN-α) treatment than HDV-1 patients^7^. The clinical features of recently identified genotypes HDV-6 through -8 are less characterized^8^.

HDV recruits HBV surface envelope proteins (HBsAg) for its progeny envelopment, which is critical for HDV propagation and transmission^4^. Similar to HBV, HBsAg-enveloped HDV virions enter hepatocytes through the interaction with the integral transmembrane protein receptor Na^+^- taurocholate co-transporting polypeptide (NTCP)^4^. NTCP not only governs HDV liver tropism, but also determines its species-specificity: only human and to a lesser extent mouse and rat NTCP homologues support HDV entry, but not pig or macaque NTCP^9–11^.

The HDV genome replicates in the nucleus using a double rolling circle mechanism, generating intermediate antigenomic and genomic viral RNA strands^12^. The mRNA transcribed from the genomic RNA encodes only one open reading frame (ORF) which leads to the translation of two proteins, the small (S-HDAg) and the large (L-HDAg) hepatitis delta antigen^13^. Editing of the amber stop codon of the S-HDAg ORF in the antigenome by adenosine deaminase acting on RNA 1 (ADAR1) elongates the reading frame by 19 to 20 codons, which drives the transition from S- to L-HDAg synthesis^14^. Within the elongated terminus of L-HDAg, a recognition site for cellular farnesyltransferase is modified and allows the switch towards virion assembly and release: while S-HDAg facilitates, the farnesylated L-HDAg inhibits HDV genome replication. L-HDAg together with the S-HDAg and the HDV genome initiates the formation of the ribonucleoprotein (RNP) complex. The farnesyl group facilitates RNP attachment to the endoplasmic reticulum membrane where HBsAg is located and new HDV progenies can be assembled^19^.

To date, fundamental aspects of HDV biology remain poorly understood, including molecular details of its life cycle, the apparent genotype-dependent disease severity, and the underlying mechanisms leading to accelerated liver dysfunction^4^. As a result, off-label peg-IFN-α was the only available treatment for HDV infections for a long time. However, it cannot be used for all patients, shows only limited efficiency, and often leads to significant side effects^4,16^. Bulevirtide (BLV, brand name Hepcludex, formerly known as Myrcludex B), has been recently granted conditional marketing authorization by the European Medicines Agency (EMA)^22^. It is based on a synthetic pre-S1-derived myristoylated peptide that specifically binds to NTCP and thus blocks HBV and HDV entry. Recent real-world studies demonstrated that BLV as a monotherapy or in combination with peg-IFN-α shows profound effects on HDV serum and liver RNA with possible curative potential in a subset of patients^19^. The anti-HDV efficacy of farnesyltransferase inhibitor Lonafarnib (LNF) which inhibits HDV progeny envelopment and therefore secretion, has been demonstrated in phase 3 clinical trial when given in combination with either ritonavir or peg-IFN- _α_20.

The gap in our understanding of HDV molecular biology and the lack of a curative treatment is partially due to the restrictions of reproducible HDV cell culture systems that resemble the physiological state of hepatocytes *in vivo*. Primary human hepatocytes (PHH) provide an attractive model for HDV *in vitro* studies. However, their restricted availability, high donor-to-donor variability, their difficulties in growing and maintaining them in culture for longer periods of time, and the limited ability to manipulate them genetically impose severe limitations for reproducible and high-throughput HDV studies^21^. Upon plating, PHH susceptibility to HDV infection drops rapidly, due to the loss of hepatocyte polarization and NTCP expression^22^. On the other hand, widely available and highly reproducible hepatoma cells can be only rendered permissive to HDV infection upon induced or ectopic NTCP expression^17^. We have recently engineered a HepG2- derived cell line that in addition to NTCP co-expresses the HBsAg, allowing us to study HDV spread^23^. However, hepatoma cells retain their transformed nature and members of the drug metabolizing CYP450 family are only expressed to low levels in these cells^24^. The human liver bipotent progenitor cell line HepaRG becomes permissive to HDV infection upon differentiation into hepatocytes^25^. While differentiated HepaRGs cells retain many characteristics of PHHs, they do not replicate all^26^.

Recently, numerous protocols have been developed to differentiate human induced or embryonic pluripotent stem cells (hiPSC/ESC) into hepatocyte-like cells (HLCs) (reviewed in^27^). HLCs resemble PHHs and better recapitulate physiological features, such as drug metabolism or innate immune response, than hepatoma cells^28^. In addition, stem cells represent a reproducible and renewable resource that can be genetically manipulated. We and others have shown that hESC- or iPSC-derived HLCs are suitable for HBV^29^, hepatitis C virus (HCV)^30^ and hepatitis E virus (HEV)^31^ studies.

Here, we show that HLCs are readily permissive for HDV infection and replication of all genotypes tested. We found that the co-infection with HBV or adeno-associated virus (AAV)- mediated transduction of HBsAg in HLCs enabled HDV progeny assembly and release. We demonstrate that HDV can spread extracellularly in HLCs which prompted us to evaluate the efficacy of available anti-HDV regimens in HLCs. Finally, by challenging the cells at different stages of HLC differentiation, we demonstrated the potential of this system to identify new host factors that either promote or restrict HDV infection.

## Results

### Hepatocyte-like cells (HLCs) are permissive for HDV infection in an NTCP-dependent manner and can be co-infected with HBV

Previous studies have demonstrated the utility of stem cell-derived HLCs for hepatotropic virus infection, including HBV (reviewed in^27^). Here, we analyzed whether HLCs are likewise permissive to HDV infection.

We first used the HDV genotype (GT) 1 strain (JC126)^32^ to infect HLCs that we differentiated from the human embryonic stem cell (hESC) line WA09 based on a previously optimized protocol^33^ (Fig. 1A). By the end of the differentiation, HLCs supported hepatocyte features including indocyanine green uptake, glycogen synthesis, and the conversion of non-fluorogenic carboxyfluorescein diacetate into fluorogenic carboxyfluorescein^33^.

**Figure 1:**
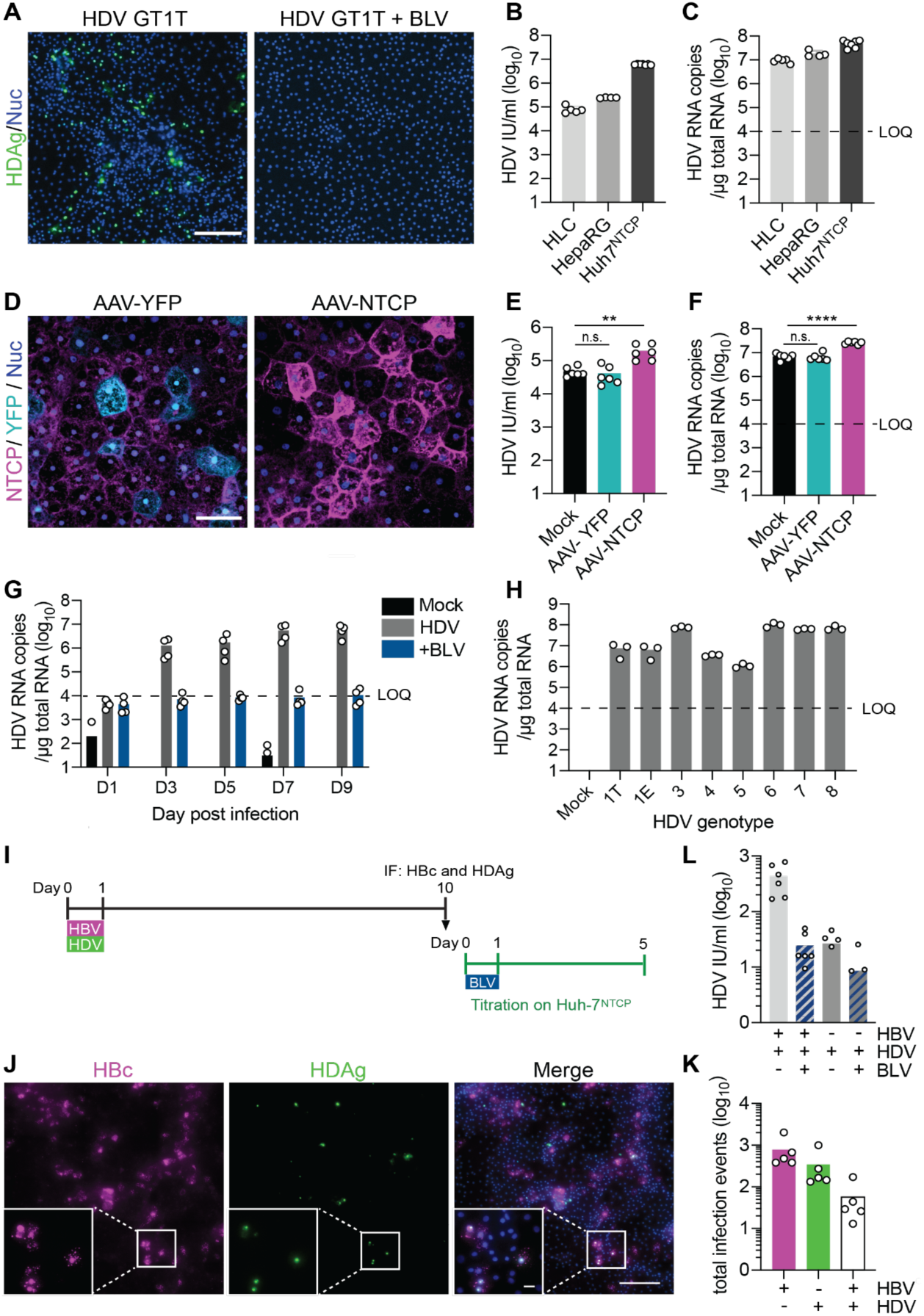
Hepatocyte-like cells (HLCs) are susceptible to HDV infection in an NTCP dependent manner. (A) HDV infection (MOI = 5) of HLCs incubated with or without 500 nM entry inhibitor bulevirtide (BLV) was assessed by immunofluorescence staining (IF) against the HDV antigen (HDAg, green) five days post- infection (p.i.). Scale bar = 200 μm. (B & C) HLCs, differentiated HepaRG, and Huh7^NTCP^ cells were infected with HDV (MOI = 5). HDV infection was analyzed by (B) counting HDAg-positive cells using CellProfiler or by (C) quantifying HDV genome copies by RT-qPCR. Dashed line: limit of quantification (LOQ); (D) HLCs were transduced with or without AAV (MOI = 10^4^ viral genomes, vg/cell) encoding for YFP or NTCP and stained with Atto-MyrB-565 (magenta) two days later. (E & F) Two days post-transduction, HLCs^YFP/NTCP^ were infected with HDV (MOI = 5 IU/cell) and analyzed by (E) counting HDAg-positive cells or (F) quantifying HDV genome copies five days p.i. Scale bar = 50 μm. Statistical analysis was performed by unpaired two-tailed Student’s t test *: *p*<0.05; **: *p* <0.01; ***: *p*<0.001; ****: *p*<0.0001; n.s.: non-significant. (G) HDV infection (MOI = 5) of HLCs incubated with or without 500 nM BLV was analyzed by quantifying HDV genome copies in HLC lysates harvested at the indicated days. (H) HLCs were infected with the indicated HDV genotype (MOI = 15 for GTs 1T, 1E, 4, 6, 7, 8; MOI = 30 for GTs 3 & 5) and five days p.i., HDV genome copies were quantified using RT-qPCR. (I) Experimental setup. HLCs were infected with HBV (MOI = 450 genome copies/cell) and HDV (MOI = 10). (J) Cells were stained against HBV core (HBc, magenta), HDAg (green), and nuclei (DAPI, blue) ten days p.i. Scale bar = 200 μm, scale bar of insets = 40 μm. (K) HBc- and HDAg-positive cells were counted using ZEN imaging software to quantify HBV and HDV single and co-infection events. Images are representative of three independent differentiations. (L) The supernatant from HBV/HDV- or HDV-infected HLCs collected on day ten p.i. was diluted 1:5 to infect Huh7^NTCP^ cells with or without 500 nM BLV. Infected Huh7^NTCP^ cells were fixed, stained for HDAg, and analyzed using CellProfiler. N = biological replicates.

**Table 1:**
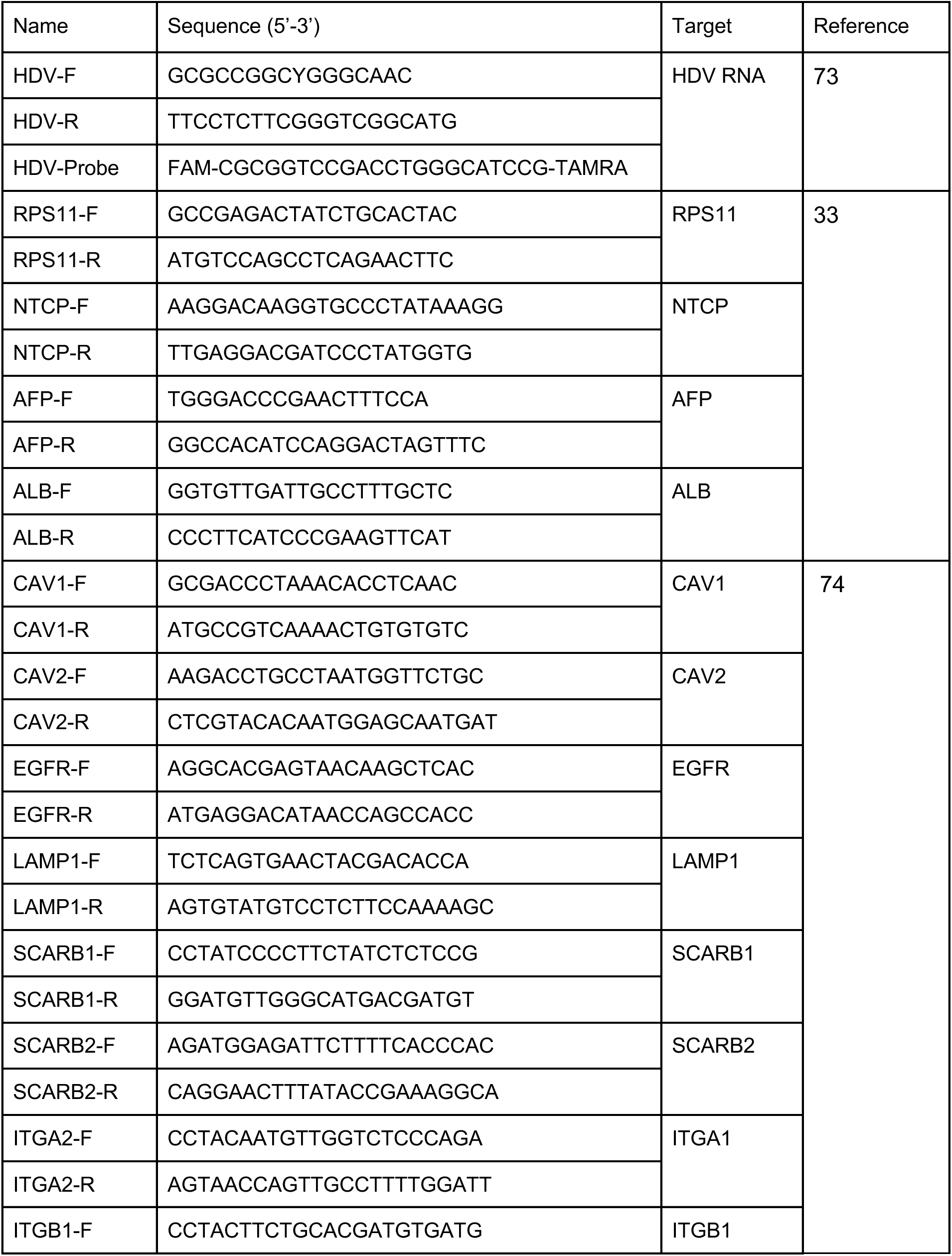
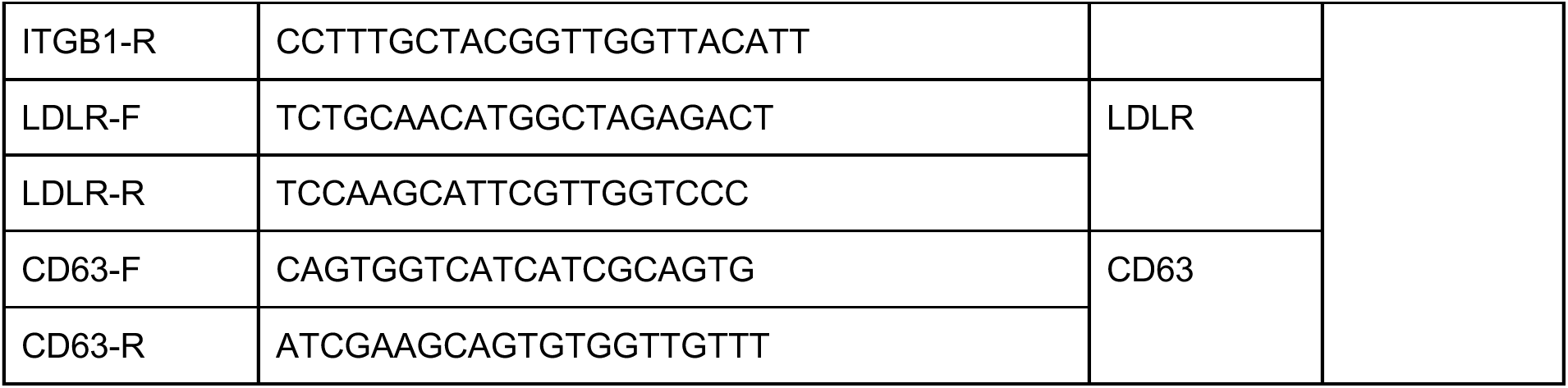
Primers used in this study.

As shown in Suppl Fig. 1A & B, we detected increasing numbers of HDAg-positive HLCs in an inoculum dose-dependent manner. HDV infection could be entirely blocked by pre-treating HLCs with the inhibitor bulevirtide (BLV) (Fig. 1A), which is based on a synthetic lipopeptide that mimics the receptor binding-site of HBV pre-S1 protein and binds to NTCP. Of note, while differentiating the cells in the final maturation medium, we observed the formation of a second, highly confluent HLC layer on top of the bottom HLC monolayer. Although we detected some HDAg-positive cells in the bottom monolayer, we found the majority of HDV infections in the top HLCs layer (Suppl Fig. 1A & Fig. 1A).

In order to evaluate the robustness of the system, we compared HDV infection of HLC with conventional HDV culture systems. When applying the same HDV inoculum, HLCs were similarly permissive to HDV infection (Fig. 1B) and replication (Fig. 1C) as differentiated HepaRG cells. However, both HLCs and HepaRG cells were much less permissive than carcinoma-derived Huh7 cells ectopically expressing human NTCP (Huh7^NTCP^, Fig. 1B & C and Suppl Fig. 1C).

Then, we assessed the possibility of further enhancing HDV infection by transducing HLCs with recombinant, NTCP-encoding adeno-associated viruses (AAV) (Fig. 1D-F). We analyzed endogenous and ectopic NTCP expression by microscopy using a fluorescently conjugated peptide binding specifically to human NTCP (Fig. 1D). Unlike hepatoma cells, for which we and others have observed a significant increase in HDV susceptibility upon NTCP replenishment^17^, ectopic NTCP expression in HLCs enhanced HDV infection by only **∼**3-fold (Fig. 1E & F). This indicated that in contrast to hepatoma cells, endogenous NTCP expression levels were not majorly rate-limiting for productive HDV infection of HLCs. Of note, we also used the hESC line RUES2 and the hiPSC line iPSC.C3A^34^ (a kind gift from Stephen Duncan, MUSC) to generate HLCs that were likewise permissive to HDV infection (data not shown), demonstrating the reproducibility of the system.

Next, we analyzed HDV replication kinetics in HLCs by quantifying HDV genome copies over time (Fig. 1G) up to 20 days post-infection (Suppl. Fig 1D). We observed a delayed onset of replication by day 3 post-infection and a decrease in replication at later time points, which is similar to HDV replication kinetics in other cell culture models, including primary human hepatocytes (PHH)^35,36^. We also infected HLCs with HDV GTs 3-8^37^ and found that HLCs were permissive for all tested genotypes (Fig. 1H and Suppl Fig. 1E).

Finally, we performed HBV genotype D and HDV GT1 co-infection of HLCs (Fig. 1I-L). Although many infected HLCs were either HBV core (HBc) or HDV antigen (HDAg) single-positive, we found ca. 25% of the infected cells to be double-positive for both HBc and HDAg, suggesting productive co-infection with both viruses (Fig. 1J & K). Upon co-infection with HBV, we detected infectious HDV progenies in the HLC culture medium showing that HLCs can fully recapitulate the entire HDV life cycle (Fig. 1L).

### AAV transduction of HLCs with HBV surface antigens allows completion of the HDV life cycle in HLCs in the absence of HBV infection

It has been experimentally shown that HBsAg derived from naturally integrated HBV DNA can support production of infectious HDV virions in the absence of active HBV replication^38,39^. Corroborating this, high HDV viremia has been detected in HDV/HBV carriers with low or undetectable HBV levels^40^, suggesting that HBV integrates are sufficient for expressing the HBV envelope proteins.

To mimic this situation, we used a previously identified AAV capsid that transduces HLCs at a high efficiency^41^ to express L-/M-/S-HBsAg in fully matured HLCs (HLCs^HBsAg^, Fig. 2A). We replaced the original promoter in the AAV transgene vector with the authentic HBV promoter in order to gain the optimal ratio between the three envelope proteins enabling particle formation, as we have shown before^23^.

**Figure 2:**
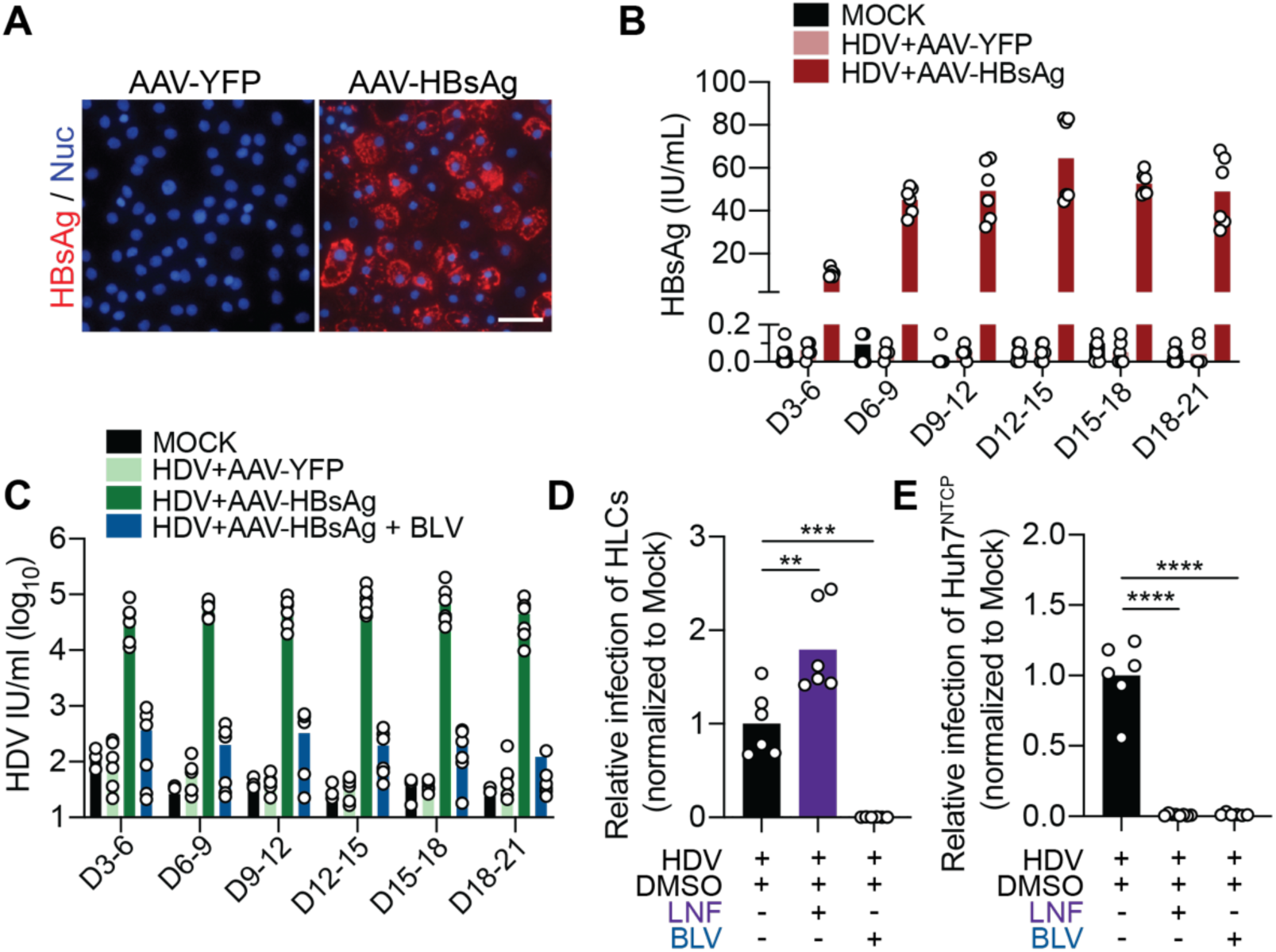
AAV transduction with HBV surface antigens allows completion of the HDV life cycle in HLCs. (A) HLCs were infected with HDV (MOI = 5) and the next day transduced with AAV-YFP or AAV-HBsAg. SN: supernatant. Nine days post-transduction, HLCs were stained for HBsAg (red) and nuclei (blue). Images are representative of two independent differentiations. Scale bars = 50 μm. (B) HBsAg was quantified in the supernatant by ELISA collected at the end of indicated time periods. (C) Progeny HDV from HLCs harvested at the indicated time points was diluted 1:5 and used to infect Huh7^NTCP^ cells with or without 500 nM BLV. Infected Huh7^NTCP^ cells were fixed and stained for HDAg to quantify HDV infections. (D) HLCs were infected with HDV (MOI = 5), transduced with AAV-HBsAg and incubated with or without 500 nM BLV (between D0-D1 p.i.) or 2 μM Lonafarnib (LNF; between D0-D5 p.i.). HDV infection was quantified by counting HDAg-positive HLCs five days p.i. (E) The supernatant from these HLCs was diluted 1:5 to infect Huh7^NTCP^ cells which were analyzed for HDV infection by HDAg staining five days p.i. N = biological replicates. Statistical analysis was performed by unpaired two-tailed Student’s t test. **: *p* <0.01; ***: *p*<0.001; ****: p<0.0001; n.s., non-significant.

Six days post-HDV-infection, we detected secreted HBsAg in the supernatant of HLCs^HBsAg^ (Fig. 2B) as well as HDV progenies, which were capable of initiating secondary infections in Huh7^NTCP^ cells (Fig. 2C). HDV-infected HLCs^HBsAg^ continued to secrete both HBsAg and infectious progenies until the end of the observation time, i.e., up to 21 days post-HDV infection.

In order to ensure that the secondary infections were not carry over events from the initial HDV inoculum, we used the HBV/HDV entry inhibitor BLV and Lonafarnib (LNF), which prevents prenylation of the C-terminal Cys211 residue in L-HDAg and thus the envelopment of infectious HDV progenies^4^. LNF increased primary HDV infection of HLCs (Fig. 2D) in agreement with previous results^23^ but it fully blocked the assembly and release of infectious HDV progenies (Fig. 2E). As BLV already very potently blocked primary HDV infection of HLCs (Fig. 2D), we did not observe any secondary infections in Huh7^NTCP^ (Fig. 2E). These data showed that HLCs can be genetically engineered to express HBsAg allowing HDV to complete its life cycle.

### HDV extracellular spread in HLCs^HBsAg^ enables the evaluation of available antiviral HDV regimens

HDV extracellular spread, which occurs efficiently *in vivo*, is difficult to replicate in currently available *in vitro* models^21,23^. Based on our observation that HDV progenies can be released from HLCs^HBsAg^, we analyzed whether they also supported HDV extracellular spread (Fig. 3). The macromolecular crowding agent polyethylene glycol 8000 (PEG) is widely used to enhance HBV and HDV infection in cell culture^21,42,43^ and we likewise used PEG 8000 for our primary HDV infections of HLCs. To confirm that HDV extracellular spread without further addition of PEG is possible in HLCs, we first confirmed that HLCs can be infected with HDV in the absence of PEG (Suppl Fig. 2A & B).

**Figure 3:**
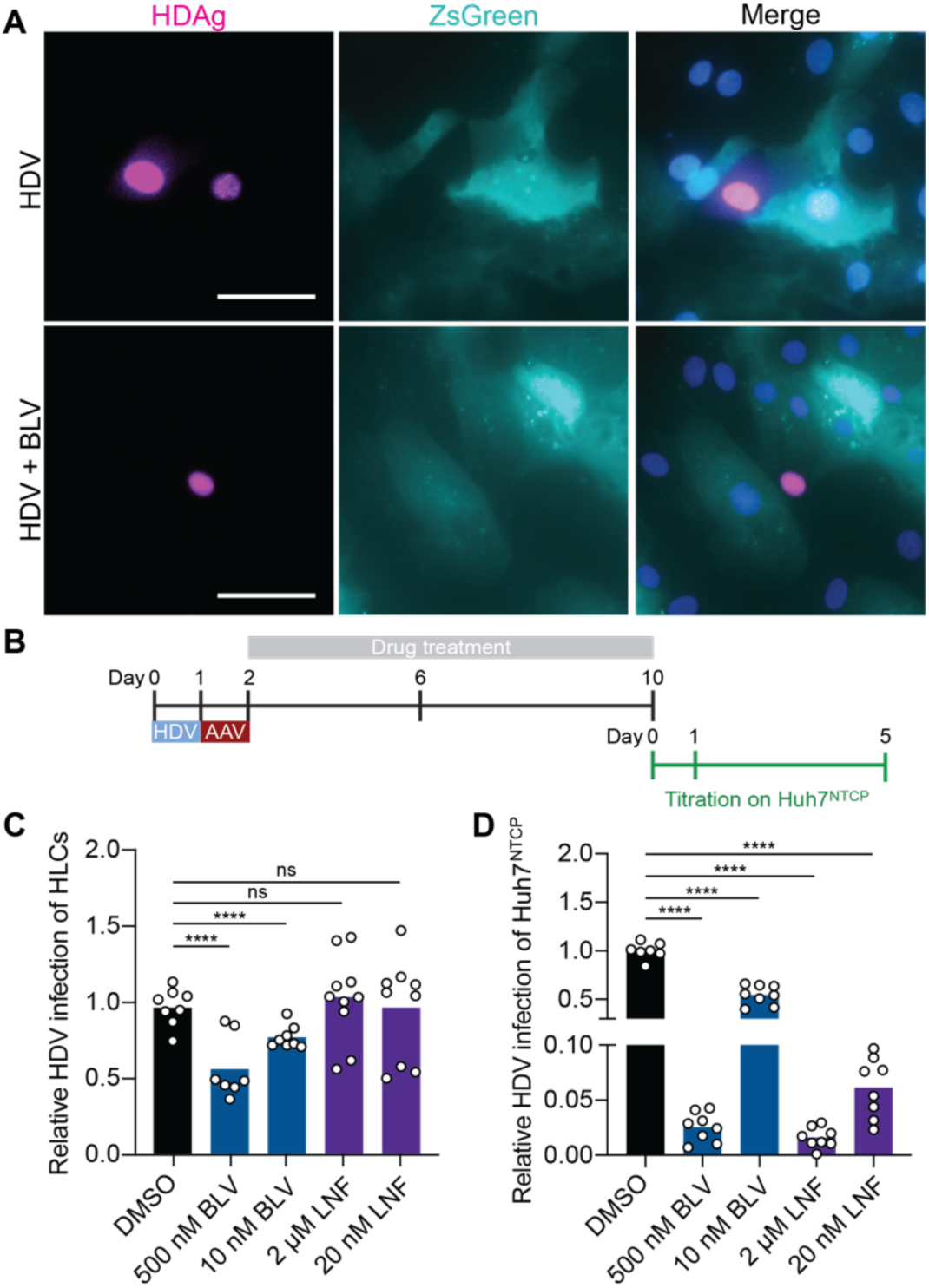
HDV spread in HBsAg-HLCs allows the evaluation of antiviral HDV regimen. (A) Wild-type HLCs were infected with HDV (MOI = 5) and the next day transduced with AAV-HBsAg. Two days post-infection, they were dissociated and co-cultured with Zs-Green expressing HLCs in the presence or absence of BLV. Eight days later, cells were fixed, stained, and imaged for HDAg (magenta), ZsGreen (cyan) and nuclei (DAPI, blue). Scale bar = 50 μm (B) Experimental setup. HLCs were infected with HDV (MOI = 5) and the next day transduced with AAV-HBsAg. After removal of the inoculum on day 2 p.i., HLCs were incubated with drugs, which were replenished every four days. Ten days p.i., HLCs were fixed to analyze HDV infections and their culture supernatant harvested for titration of HDV progenies on Huh7^NTCP^ cells. (C) Relative HDV infection events normalized to vehicle DMSO-treated cells were quantified by counting HDAg-positive HLCs 10 days p.i. (D) The supernatant from HLCs was diluted 1:5 to infect Huh7^NTCP^ cells, which were then analyzed for HDV infection by HDAg staining five days p.i. N = biological replicates. Statistical analysis was performed by one-way ANOVA **: *p* <0.01; ***: *p*<0.001; ****: p<0.0001; n.s., non-significant.

Next, we transduced WA09 cells at their pluripotent stem cell stage with lentiviruses to express ZsGreen and differentiated them to HLCs^ZsGreen^. In parallel, we differentiated wild-type WA09 cell and infected them with HDV and transduced them with AAVs to express HBsAg. We then detached both HLC types and mixed the “recipient” HLCs^ZsGreen^ with “donor” HLCs^HDV/HBsAg^. Eight days later, we analyzed co-cultured donor and recipient HLCs for HDV infection by staining against HDAg. As shown in Fig. 3A, we found HDAg-positive HLCs^ZsGreen^ in the vicinity of HDV- infected donor HLCs. The treatment with BLV completely prevented infections of HLCs^ZsGreen^, showing unequivocally that HDV extracellular spread had occurred in our HLC culture.

This prompted us to test the impact of available anti-HDV treatments on HDV spread in HLCs (Fig. 3B). First, we infected HLCs with HDV to allow primary infection. The next day, we transduced them with AAV-HBsAg. On the third day, we added the drugs to evaluate their impact on the secondary infections, and accordingly, extracellular spread only. Since HLCs similar to PHHs do not proliferate^44^, we could exclude cell-division mediated spread.

Ten days post-infection, we observed that the treatment with a relatively high dose of 500 nM BLV decreased the total HDV infection events compared to non-treated HLCs by roughly 50% (Fig. 3C). In a dose-dependent manner, 10 nM BLV decreased total HDV infections by only ∼25%. Interestingly, the treatment with LNF, which prevents HDV assembly, did not have any impact on the total number of HDV infections by day 10, but showed an effect when using the supernatant to titer infectious progenies on Huh7^NTCP^ cells (Fig. 3D). This may be due to the observation that LNF enhances primary infections (Fig. 2D) which is in agreement with reports by others^23,45^. Of note, the high dose of BLV completely abrogated secondary infection of Huh7^NTCP^ cells (Fig. 3D), although some progenies must have been released from primary infected HLCs. This suggested carry-over of BLV in the inoculum and highlights the advantage of using a culture system that supports authentic and extracellular HDV spread.

### HDV susceptibility along HLC differentiation

We finally wanted to determine the stage along the differentiation process at which HLCs became susceptible to HDV infection. HLC differentiation is based on a five-step protocol mimicking liver development with each step involving the exposure of the cells to different culture media and growth factors: from hESC to definitive endoderm (DE), followed by hepatic specification, then immature, and finally mature hepatocytes (Fig. 4A). We monitored the differentiation progress by analyzing the expression of proteins (Fig. 4A) and transcript levels (Fig. 4B) of the immature hepatocyte marker alphafetoprotein (AFP), the mature hepatocyte marker albumin (ALB), and NTCP. As soon as we cultured DE cells in the medium containing hepatocyte growth factor to initiate hepatic specification, we observed a steep increase in AFP, ALB, and NTCP expression levels (Fig. 4B). While AFP levels reached a plateau by day 13, ALB expression only peaked by day 19 in the last culture medium, when the cells reached their final maturation stage. Interestingly, NTCP was already expressed on the cell surface during the hepatic specification stage but slightly decreased during the final maturation step.

**Figure 4:**
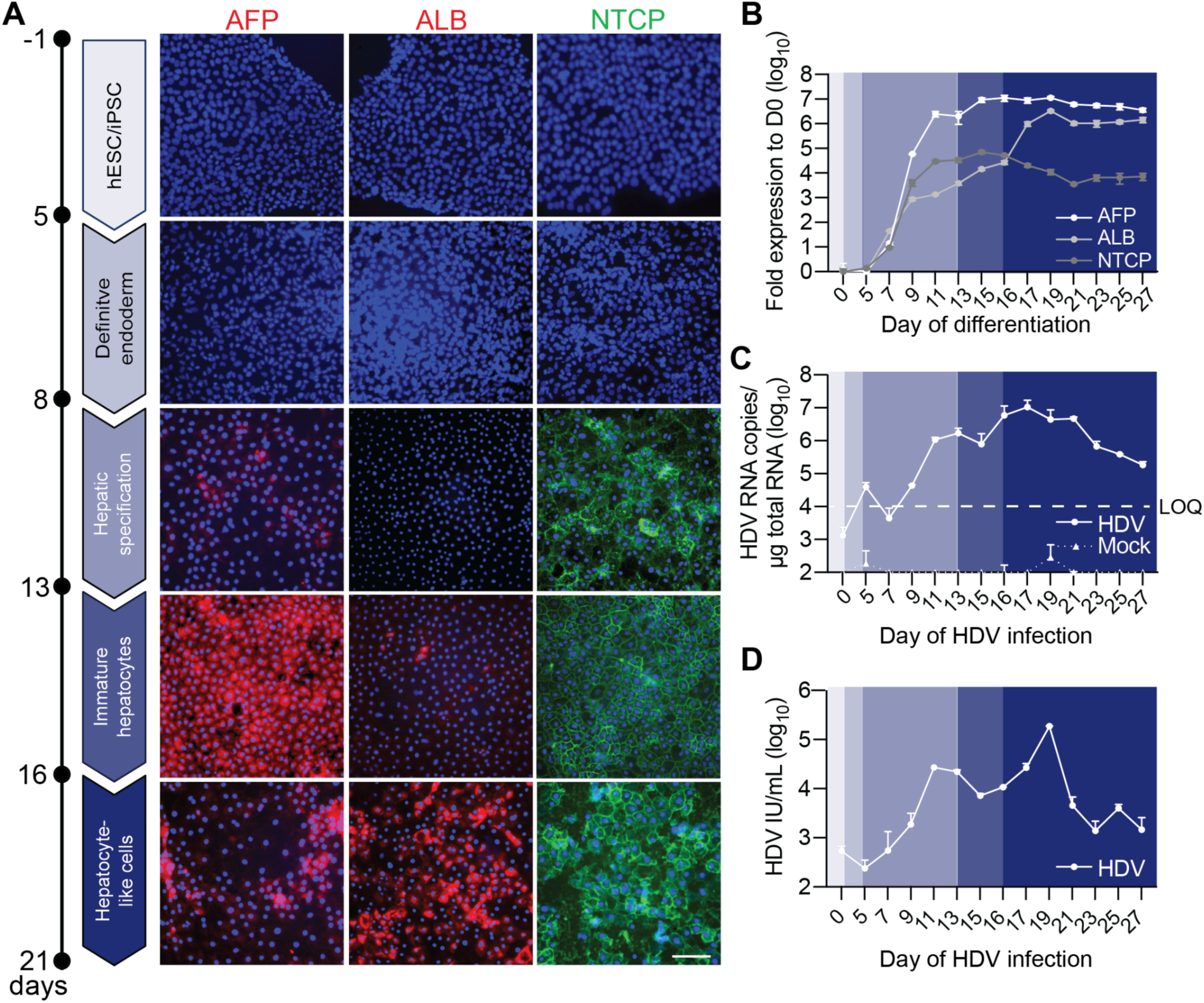
HDV susceptibility along stem cell differentiation to hepatocyte-like cells. (A) Immunofluorescent images of cells stained against the nuclei (DAPI, blue) and with antibodies against alphafetoprotein (AFP, red), albumin (ALB, red) or Atto-MyrB^488^ (NTCP, green) at the following stages during HLC differentiation: hPSCs, definitive endoderm, hepatic specification, immature, and mature hepatocyte-like cells. Images are representative of two independent differentiations. Scale bar = 100 μm. (B) Cells were harvested for analyzing ALB, AFP, NTCP expression using RT-qPCR at the indicated day of the differentiation protocol. (C & D) Cells were infected with HDV at the indicated day of the differentiation protocol and harvested five days p.i. HDV infection was analyzed by quantifying (C) HDV genome copies and (D) HDAg-positive cells using CellProfiler. Dashed line: LOQ. Results represent the mean ± SD of N = 2.

Then, we infected the cells at either stem cell level (day 0), DE stage (day 5), or every second day during the hepatocyte differentiation with HDV. 5 days post-infection we analyzed HDV infections by quantifying HDV RNA copy numbers (Fig. 4C) and HDAg-positive cells (Fig. 4D) five days post-infection. Neither hESC nor DE cells were susceptible to HDV infection (Fig. 4C & D). Interestingly, when bypassing the cell entry step by delivering the HDV genome via transfection, we found that hESCs were already capable of replicating HDV, as evidenced by the detection of L-antigen in transfected cells (Suppl Fig. 3A).

We found that hepatic progenitors became susceptible to HDV infection as early as day 11 after starting the differentiation protocol, likely governed by the expression of NTCP. Interestingly, immature hepatocytes at day 15 were less susceptible to HDV infection than the progenitors, although they expressed NTCP on their surface. Then, we observed a second peak of HDV infection, when the cells acquired a fully matured hepatocyte profile, as evidenced by the peak in ALB expression at day 19 (Fig. 4B). Remarkably, NTCP expression was not further enhanced at this point in time, indicating that other factors restricted HDV at the immature state. To identify the exact day, we further refined and infected the cells with HDV every day between day 17 to 19 (Suppl Fig. 3B & C) of the differentiation protocol. We identified day 18 as the differentiation day when cells became the most susceptible to HDV infection, roughly 2 days after we switched to the final maturation medium. From day 19 on, HLCs became less susceptible to HDV infection, likely due to a further decrease in NTCP expression (Fig. 4B) from the mature HLC stage on.

### Differential gene expression analysis reveals potential co- and restriction factors of HDV infection in mature HLCs

The enhanced permissiveness observed in matured HLCs by day 18 could be governed by the upregulation of a host co-factor or the downregulation of a host restriction factor for HDV infection. Thus, we compared the transcriptome profile of the cells between day 17 and day 18 (Fig. 5). Whole-transcriptome expression profiling and gene ontology (GO) term enrichment analysis revealed up- or downregulation of genes enriched in many pathways, such as liver development and regeneration (Fig. 5A). In agreement, individual markers of mature hepatocytes such as ALB and PROX1 were upregulated, while markers of immature hepatocytes, such as AFP and the stem cell marker SOX9, were downregulated by day 18 (Fig. 5B). FRZB is a negative regulator of hepatocyte differentiation and was likewise downregulated by day 18.

**Figure 5:**
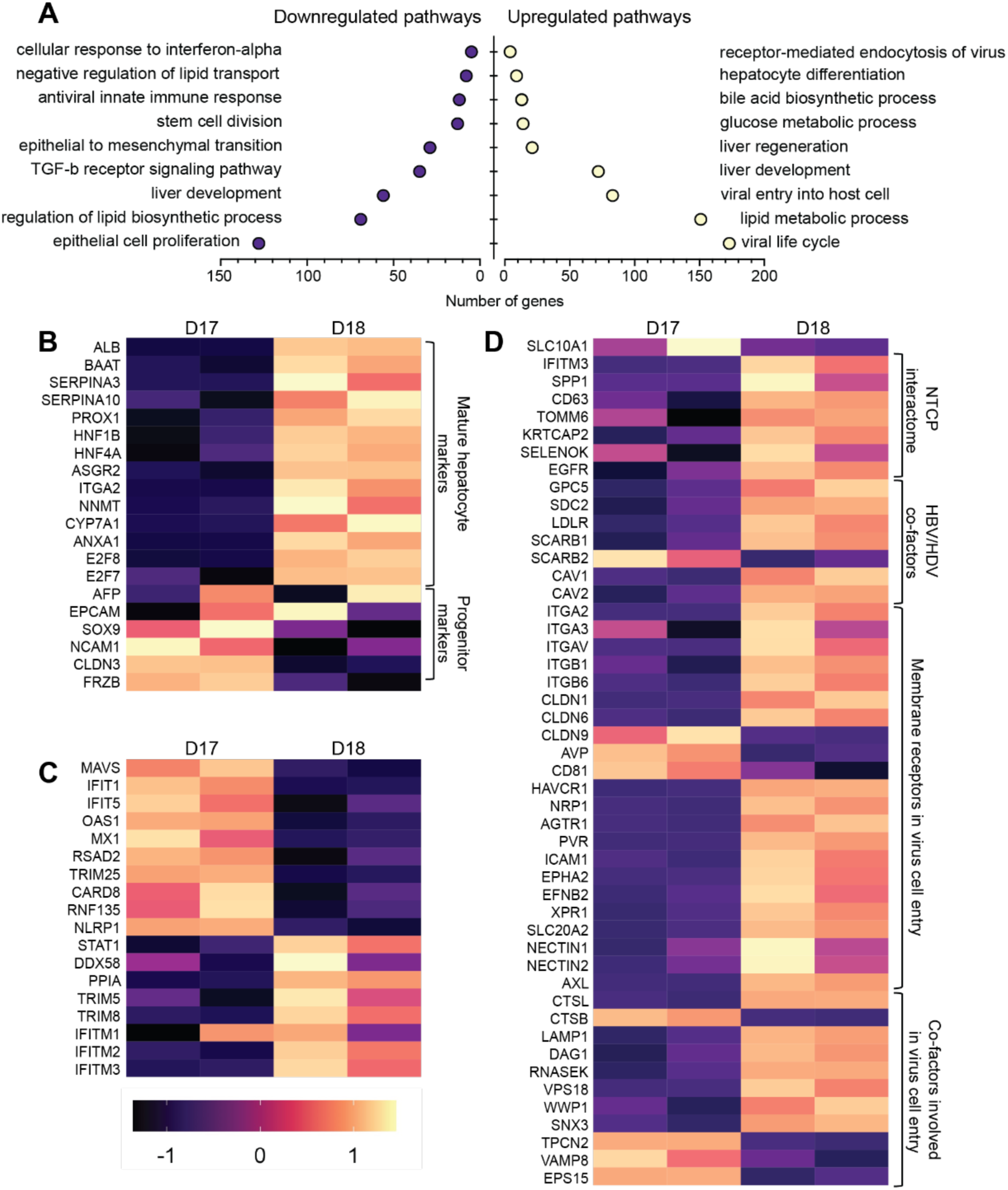
Differential gene expression analysis reveals upregulation of HBV/HDV entry factors in mature HLCs. Total RNA was extracted from HLCs at either day 17 or 18 during the differentiation protocol and subjected to whole-transcriptome expression profiling and gene ontology (GO) term enrichment analysis. (A) GO term enrichment analysis of biological pathways for up- and down-regulated genes between HLCs at day 17 and 18. Differentially expressed genes (p value <0.05) were significantly enriched in this GO term. (B-D) Heatmap of *Z* score-normalized counts per million (CPM) values for (B) hepatocyte markers, (C) innate immune genes, and (D) virus entry factors. N = biological replicates.

We previously showed that stem cells express an intrinsic and protective subset of antiviral genes (interferon stimulated genes, ISGs) and that they become downregulated throughout the differentiation^46^. In return, differentiated cells become more IFN-signaling-competent. In agreement, we observed the downregulation of a subset of ISGs (Fig. 5C). Other factors such as DDX58 and STAT1, which are critical for IFN induction and signaling, respectively (Fig. 5C), and which rather play an antiviral role in mature hepatocytes, were upregulated.

Furthermore, the GO analysis revealed the upregulation of many genes that have been previously described to be involved in the entry process of other viruses, including HBV (Fig. 3D). Notably, genes encoding proteins that were found to interact with NTCP^47^ as well as previously described HBV and HDV co-entry factors were upregulated in mature HLCs by day 18. In addition, numerous membrane receptors and co-factors involved in general viral entry were likewise upregulated. We validated by RT-qPCR the upregulation of HBV entry co-factors along HLC differentiation (Suppl Fig. 5A), such as caveolins^48^ and LAMP1 as well as previously described membrane receptors (Suppl Fig. 5B), such as EGFR^49^, SCARB1^50^, and LDLR^51^.

### CD63 is a potential co-factor of HDV entry and could be rate-limiting for infection of immature hepatocytes

In order to identify a potential host factor that was expressed in mature HLCs and responsible for enhanced HDV infection, we performed a small siRNA screen in Huh7^NTCP^ cells. We selected siRNAs that target genes encoding membrane receptors and co-factors of viral entry and that were upregulated in mature HLCs as revealed by the transcriptome analysis (Fig. 5D). As positive controls, we also employed siRNAs targeting NTCP as well as the HDAg-coding region in the HDV RNA genome.

As shown in Fig. 6A, the downregulation of several genes reduced HDV infection by up to 80% compared to the transfection with a non-target (nt) control siRNA. We selected the top four candidate genes for further validation (Fig. 6B): the poliovirus receptor (PVR), CD63, as well as integrins beta 1 (ITGB1) and alpha 2 which can form a functional ITGB1/A2 heterodimer. We confirmed the downregulation of these genes by western blot analysis (Suppl Fig. 5A) and their impact on HDV infection of Huh7^NTCP^ cells (Fig. 6B). However, the downregulation of PVR and ITGA2 had a pronounced effect on cell attachment and cell viability (Fig. 6C). Since the downregulation of CD63 decreased HDV infection by up to 50% in the absence of any effect on either surface NTCP expression (Suppl Fig. 5B) or cell viability (Fig. 6C), we decided to follow up on this host factor.

**Figure 6:**
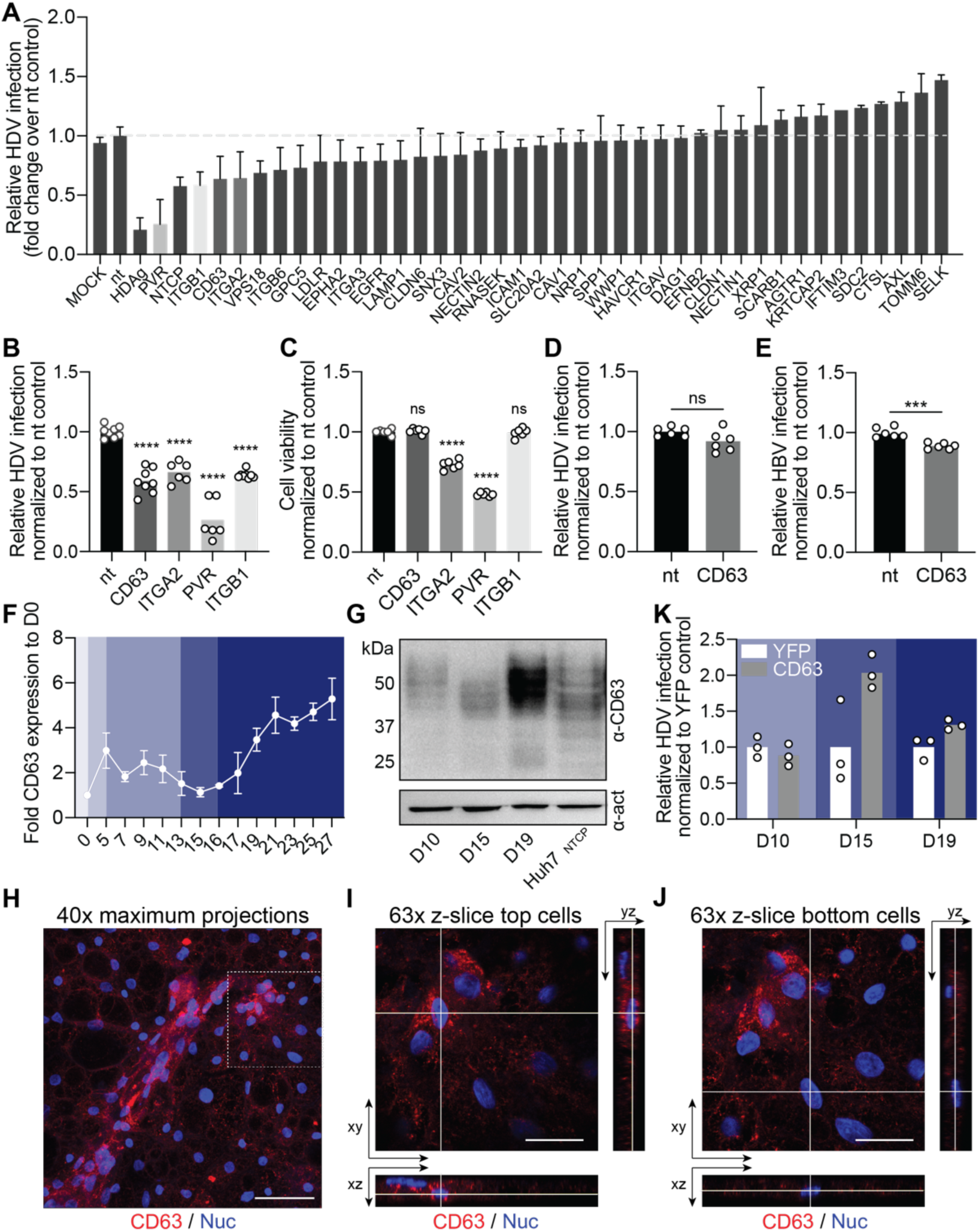
siRNA screen reveals CD63 to be potential co-factor of HBV/HDV entry which could be rate-limiting for infection of immature hepatocytes. (A) siRNA screen of potential HDV host factors. Huh7^NTCP^ cells were transfected with 50 nM on-target pool siRNAs directed against indicated genes and 24 h later infected with HDV (MOI = 1). Relative HDV infection was normalized to non-target (nt) siRNA transfection and quantified by counting HDAg-positive cells five days p.i. (B & C) Four genes were selected and confirmed by separate siRNA transfection into Huh7^NTCP^ cells and analyzed for (B) HDV infection and (C) cell toxicity. Statistical analysis was performed by one- way ANOVA **: *p* <0.01; ***: *p*<0.001; ****: p<0.0001; n.s., non-significant. (D) Huh7^NTCP^ cells were infected with HDV (MOI =1) and 24 h later transfected with nt- and CD63-siRNAs. HDV infections were quantified by counting HDAg-positive cells five days p.i. Statistical analysis was performed by unpaired two-tailed Student’s t test. **: *p* <0.01; ***: *p*<0.001; ****: p<0.0001; n.s., non-significant. (E) HepG2^NTCP^ cells were transfected with nt- and CD63-siRNAs and 48 h later, infected with HBV (MOI = 300 genome copies/cell). HBV infections were quantified by counting HBV core-positive cells 10 days p.i. Statistical analysis was performed by unpaired two-tailed Student’s t test. **: *p* <0.01; ***: *p*<0.001; ****: p<0.0001; n.s., non-significant. (F) HLCs were harvested for analyzing CD63 expression using RT-qPCR at the indicated day of the differentiation protocol. (G) Western blot analysis of Huh7^NTCP^ cells as control and HLC cell lysates harvested at indicated day of the differentiation protocol, for CD63 and β-actin (act) expression using respective antibodies. (H-I) HLCs at their fully matured stage were fixed, stained for CD63 (red) and the nucleus (blue) and imaged at the Airyscan confocal microscope. (H) Maximum intensity projections on 40x tile image stacks showing both, top and bottom HLCs. Scale bar = 50 μM. Single z-slice and orthogonal xz and yz views of (I) top HLCs or (J) bottom HLCs. Scale bar = 20 μM. (K) Cells at indicated day of the differentiation protocol were transduced with AAV-YFP or AAV-CD63 and two days later, infected with HDV. Relative HDV infections were quantified by counting HDAg-positive cells five days p.i. N = biological replicates.

Next, we wanted to study whether CD63 plays a role in either the early steps of the HDV life cycle, including cell entry, or at later steps, such as genome replication. Thus, we also delivered the CD63 targeting siRNA one day after HDV infection of Huh7^NTCP^ cells (Fig. 6D). Only when CD63 was downregulated before (Fig. 6A & B) but not after HDV infection, we observed a significant reduction, suggesting that CD63 plays a role during the early steps of HDV infection, but not replication. Since HBV shares the envelope glycoproteins with HDV, we also wanted to analyze whether CD63 could play a role during HBV infection. As shown in Fig. 6E, although CD63 downregulation had a significant effect on HBV infection, it affected it to a much lower extent than HDV infection.

When analyzing CD63 expression along HLC differentiation, we found a distinct upregulation in mature HLCs as compared to immature HLCs, on the transcript and protein level (Fig. 6F & G). Interestingly, we also found that CD63 appeared to be less glycosylated in immature hepatocytes as compared to mature hepatocytes (Fig. 6G), mirroring observations made in immature and mature dendritic cells by others^52^. As described above and as shown in Fig. 1A and Suppl Fig. 1A, we observed throughout our experiments that HDV seemed to preferentially infect the highly confluent, second layer of HLCs on top of the monolayer of HLCs at the bottom. By confocal microscopy analysis, we indeed found that CD63 was more highly expressed in the top HLCs layer (Fig. 6H & I) as compared to the lower HLC level (Fig. 6H & J).

We then wanted to rescue CD63 expression at the different hepatocyte maturation stages. To this end, we AAV-transduced the cells at days 8, 13, and 17 of the differentiation protocol and infected them with HDV at each time point two days later (Fig. 6K). Ectopic CD63 expression led to a stronger, at least 2-fold increased HDV infection in immature hepatocytes but only to a 1.2- fold increase in mature hepatocytes, showing that CD63 could be a rate-limiting factor for HDV- infection of immature hepatocytes. Future studies shall provide a more mechanistic understanding on how CD63 may be involved in HDV entry. Here, we provided a guideline of how stem cell differentiation culture systems can be used as a platform for the identification of novel co-factors for virus infection.

## Discussion

Historically, studies of human hepatotropic viruses were hampered by the lack of physiologically relevant hepatic cell culture models. Likewise, for HDV studies, the scarceness of reproducible cell culture systems that resemble the physiological status of hepatocytes *in vivo,* has hampered our progress in understanding important aspects of HDV biology. Previous studies by us and others have shown the advantages of hPSC-derived HLCs for HBV, HCV, and HEV investigations^29,31,53^. Here, we demonstrated the applicability of HLCs for HDV studies.

### HLCs to study the entire life cycle of HDV and HBV/HDV co-infections

We found that HLCs were readily permissive for HDV and HBV infection and that the ectopic NTCP expression under a foreign promoter did not dramatically increase HDV infection. These results are in stark contrast to hepatoma cells^17^ and highlight the authenticity of HLCs. Thus, HLCs enable HDV infection studies under the physiological regulation of NTCP expression, which can depend on bile acid concentrations, inflammatory cytokines, and others (reviewed in^54^).

During the work on this manuscript, a study by Lange and colleagues^45^ used HLCs to study the innate immune responses to HDV mono-infection. Here, in contrast, we studied HDV biology in the context of its helper virus. By either co-infecting with HBV or by genetically manipulating HLCs to express HBsAg, we detected infectious HDV progenies in the HLC culture medium showing that HLCs can recapitulate the entire HDV life cycle and even extracellular HDV spread. Both *de novo* infection and cell division-mediated spread contributes to HDV persistence in CHD patients^55^. We have previously shown that HDV spreads in hepatoma cells upon cell division and that it was highly dependent on the innate immune competence of the respective cell line used^56^. However, HDV extracellular spread is difficult to replicate in available *in vitro* models. On the one hand, rapid de-differentiation and rapid decrease of NTCP expression after seeding heavily restrict the use of PHHs for HDV spread studies^22^. On the other hand, hepatoma cells ectopically expressing NTCP and HBsAg only poorly support extracellular spread, as shown in our recent study^23^ Here, we showed that HLCs support efficient extracellular HDV spread and thus provide a physiologically relevant, reproducible, and non-proliferative cell model to study the underlying determinants.

### HBsAg-HLCs for anti-HDV treatment evaluation

HLCs express members of the CYP450 family and their drug responsiveness correlates with those of PHHs^57^. Therefore, numerous previous studies have proposed HLCs as a platform for drug toxicity^58^ and drug evaluation studies^28^. In addition, owing to the self-renewal of stem cells, HLCs can be easily upscaled and used for high-throughput drug screens^59^. Finally, patient-specific iPSCs can be generated from clinically relevant individuals to create personalized disease and drug evaluation models.

Since HLCs^HBsAg^ supported extracellular HDV spread, we used them to test the currently available drugs specifically developed to treat HDV infections. First, we confirmed that BLV completely blocked HDV entry into HLCs. In contrast, the application of LNF led to more detectable primary HDV infections, which may be related to intra-hepatocellular accumulation of non-farnesylated and therefore non-inhibitory L-HDAg^23^. As a result, we observed no significant variation in total HDV infections after the treatment of LNF in our HLC spread assay. Only when titrating HDV progenies separately and analyzing secondary infections, we could observe the effect of LNF. Our findings highlight the advantage of studying the potency of drugs in a system that faithfully recapitulates spread, especially since the currently available and specific anti-HDV regimens target virus entry and assembly.

### Challenging HLCs along their differentiation revealed CD63 as a potential HDV/HBV entry factor

To our surprise, NTCP was expressed very early during HLC differentiation which mimics liver development. This early expression of NTCP rendered hepatic progenitors susceptible to HDV infection, aligning with previous studies that showed non-hepatic cell lines becoming susceptible to HDV infection upon ectopic NTCP expression^11^. In agreement, we found stem cells to be already capable of replicating the HDV genome when delivered through transfection, bypassing the entry step, which is consistent with the findings by Lange et al.^45^

Surprisingly, we discovered that fully matured HLCs at day 18 of differentiation exhibited a higher susceptibility to HDV infection compared to less mature cells. This suggested the presence of one or several hepatocyte-specific co-factors that facilitated HDV entry or replication. We ruled out NTCP as the responsible determinant, as its expression was slightly decreased in fully matured hepatocytes.

Our subsequent analysis using a small siRNA screen targeting selected genes from the transcriptome analysis, revealed CD63 as a potential HBV and HDV entry factor. CD63 seemed to be rate-limiting for HDV infection of immature hepatocytes which was alleviated by ectopic CD63 expression. While CD63 is most prominently known for its role in exosomal egress and has been reported to be involved in HBV assembly and egress^60^, it is also a critical component of late endosomes and facilitates vesicular trafficking through endosomal pathways^61^. As such, it has been shown to be involved in the entry of other viruses^62,63^.

It is important to note that CD63 is likely not the only factor governing enhanced infection of mature HLCs, and future research should aim to identify and confirm such other factors. In addition, further mechanistic studies will be needed to confirm and elucidate the role of CD63 in HBV/HDV entry. In this study, we demonstrated how challenging cells along the stem cell differentiation process can be a dynamic platform for discovering new host factors for virus infection.

### Limitations of using HLCs for HDV infection studies

While we have shown that HLCs support HDV infection, their susceptibility remained inferior to the infection of hepatoma cells overexpressing NTCP, potentially due to their immature nature. In addition, we observed that HDV preferentially infected the top layer of HLCs but not the bottom layer. While we found a potential correlative expression with CD63, future studies shall identify other critical co-factors for HDV infection being absent or potential restriction factors being present in the different HLC populations. Comparing the genetic landscape between the two populations could potentially lead to the identification of such factors. These factors could then be genetically modified by lentiviral or AAV transduction, as we have done in this study, to yield highly permissive HLCs.

Generally, stem cell culture and their differentiation remain expensive and time-consuming. In the absence of a deep understanding of the molecular mechanism of liver development and regeneration^64,65^, a broad range of HLC differentiation protocols have been published for different applications. A better understanding of liver development is needed and shall lead to the development of more robust and potentially commercially available HLC differentiation kits in the future. This should make the system available to all researchers in the field with the overarching goal of advancing molecular HDV studies and developing curative and alternative therapies for chronic HDV patients.

## Materials and methods

### Reagents and antibodies

The following antibodies were used for immunofluorescence staining or western blot analysis: monoclonal mouse/human anti-HDAg^37^ (1:3000, FD3A7; available through Absolute Antibody) and rabbit anti-HBcAg antibody^66^ (1:1000, H363) were generated in-house. Human anti-HBsAg (1:1000, HBc34) was a kind gift from Davide Corti, Humabs BioMed. Rabbit anti-FoxA2 (1:400) was purchased from Cell Signaling, mouse anti-AFP (1:1000) from Sigma-Aldrich, mouse anti-ALB (1:1000) from Cedarlane and mouse anti-CD63 from SANTA CRUZ BIOTECHNOLOGY (1:400). Alexa Fluor 488/568 anti-mouse (1:1000), Alexa Fluor 488 anti-human (1:1000), and Alexa Fluor 488/568 anti-rabbit (1:1000) antibodies were purchased from Invitrogen. Antibodies for western blot: mouse anti-CD63 (1:1000) was purchased from Invitrogen, rabbit anti-PVR (1:1000) from Sigma-Aldrich, mouse anti-ITGβ1 (1:1000) from Santa Cruz and rabbit anti-ITGα2 (1:1000) from Abcam. Lonafarnib was purchased from Selleckchem.

### Standard cell culture

The human hepatoma cell line Huh7^NTCP^ and HepG2^NTCP^ ectopically expressing human NTCP was generated previously^66^. Huh7^NTCP^ and HepG2^NTCP^cells were cultured in Dulbecco’s modified Eagle medium (DMEM, Gibco) supplemented with 10% advanced fetal bovine serum (FBS, Capricorn) and 2 µg/mL puromycin (InvivoGen). For AAV production, HEK-293T cells were cultured in DMEM (Gibco), supplemented with 10% fetal bovine serum and 1% penicillin-streptomycin (Gibco).

### Generation of human pluripotent stem cell-derived hepatocyte-like cells (HLCs)

The use of human embryonic stem cells (hESC) for this work was approved by the German Central Ethics Committee for Stem Cell Research (Robert Koch Institute, AZ: 3.04.02/0179). The hESC line WA09 (WiCell) was cultured in mTeSR1 medium (STEMCELL Technologies) on Matrigel (Corning) coated plates. WA09 cells were differentiated to definitive endoderm (DE) using the STEMdiff^TM^ Definitive Endoderm Kit (STEMCELL Technologies) according to the manufacturer’s protocol. To induce hepatic differentiation, DE cells were differentiated in the basal medium consisting of CTS^TM^ KnockOut^TM^ DMEM/F12 (Gibco), 10% KnockOut Serum Replacement (KOSR, Gibco), 1% MEM solution of non-essential amino acids (NEAA, Gibco), 1% GlutaMAX supplement (Gibco), and 1% penicillin-streptomycin (Gibco), supplemented with human hepatocyte growth factor (HGF, Prepotech), DMSO (Sigma-Aldrich), and dexamethasone (Sigma) as previously described^33^. For final maturation, HLCs were cultured in the Hepatocyte Culture Medium BulletKit^TM^ (HCM, Lonza) supplemented with 20 ng/mL of recombinant human oncostatin M (OSM, R&D systems). For the co-culture experiment, WA09 cells were transduced with lentivirus to express ZsGreen and selected with 1 μg/mL puromycin as described^67^ prior to HLC differentiation.

### HBV and HDV production and infection

HBV (GT D) virus was produced from the HepF19 cell line as previously described^68^. HDV virus was produced as previously described^23^. In brief, virus was collected from supernatants from Huh7 cells co-transfected with plasmids pJC126^32^ (HDV GT1, kindly provided by John Taylor, Fox Chase Cancer Center) and pT7HB2.7^69^ (hepatitis B virus genotype D envelope proteins, kindly provided by Camille Sureau, Institut National de la Transfusion Sanguine; accession number: MN645906) and purified by heparin affinity chromatography. HDV virus stocks of GT1 Ethiopia and 3-8 were produced as previously described^37^.

HLCs were infected at day 18 of the differentiation protocol at an MOI of 5 infectious units (IU) in the presence of 4% polyethylene glycol 8000 (PEG, Sigma-Aldrich) and 1.5% DMSO (Sigma-Aldrich). 16 to 24 h later, the inoculum was removed and HLCs were washed twice with Dulbecco′s Phosphate Buffered Saline (DPBS) before replenishing with fresh culture medium supplemented with 1.5% DMSO. Medium was exchanged every two days until the end of the experiment.

For the secondary infection, 1x10^5^ Huh7^NTCP^ cells were seeded in 24-well plates and inoculated with approximately one fifth of the culture supernatant from HDV-infected HLCs in DMEM containing 4% PEG and 2% DMSO (Carl Roth) after 24h. 16 to 24 h post-infection, cells were washed twice with PBS and replenished with fresh DMEM supplemented with 2% DMSO. Medium was exchanged every two days until the end of the experiment.

### Adeno-associated virus (AAV) production and transduction

The hNTCP or CD63 gene was cloned into a self-complementary AAV vector (pscAAV-CMV-EYFP-BGHpolyA^70^). The HB2.7 subgenomic fragment encoding the L-/M-/S-HBsAg of HBV genotype D was cloned into a single-stranded AAV vector (pSSV9-AAV-CMV-EYFP- BGHpolyA)^71^. Recombinant AAVs of serotype 6 were produced and iodixanol-purified as described previously^72^. HLCs were transduced with AAVs two days prior to HDV infection (to express NTCP or CD63) or one day after HDV infection (to express HBsAg). 24 h post-transduction, the inoculum was removed, HLCs were washed with DPBS, and replenished with fresh culture medium.

### Statistics

Graphs and statistical analyses were performed using GraphPad PRISM 8. In all figures where p-values were calculated, the corresponding statistical test is listed in the figure legend.

For further details regarding the materials and methods used, please refer to supplementary information.

## Abbreviations

AAV: Adeno-associated virus
ADAR1: adenosine deaminase acting on RNA 1
AFP: alphafetoprotein
ALB: albumin
BLV: Bulevirtide
CHD: chronic hepatitis D
DE: definitive endoderm
DPBS: Dulbecco′s phosphate buffered saline
EMA: European Medicines Agency
GT: genotype
HBV: hepatitis B virus
HBc: HBV core antigen
HBsAg: HBV surface protein
HCV: hepatitis C virus
HDV: hepatitis D virus
HDAg: hepatitis delta antigen
hESC: human embryonic stem cell
HEV: hepatitis E virus
HLCs: hepatocyte-like cells
hiPSC: human induced pluripotent stem cell
ISGs: interferon stimulated genes
LNF: lonafarnib
NTCP: Na^+^-taurocholate co- transporting polypeptide
pegIFN-α: pegylated interferon alpha
PHH: primary human hepatocytes
p.i: post infection
RNP: ribonucleoprotein.

## Acknowledgements

The authors gratefully acknowledge Dr. John Taylor and Dr. Camille Sureau for generously sharing plasmids. We acknowledge Dr. Vibor Laketa, head of the Infectious Diseases Imaging Platform (IDIP) at the University Hospital Heidelberg for expert support. We thank Andrew Freistaedter and Franziska Schlund for excellent technical support. We thank Dr. Yi Ni for valuable and helpful discussions.

## Authors contributions

Conceptualization, H.C., S.U., and V.L.D.T.; Methodology, H.C., and V.L.D.T.; Investigation, H.C., B.Q., A.P., L.M., J.H., R.M.F., F.A.L., Z.Z., and D.G.; Software, H.C.; Data analysis, H.C., B.Q., X.W., D.G., S.U., and V.L.D.T.; Writing-original draft, H.C. and V.L.D.T.; Final draft, H.C., S.U., and V.L.D.T.; Supervision, S.U. and V.L.D.T.; Funding, S.U. and V.L.D.T.

## Data and materials availability

All data are available in the main text or the supplementary materials.

