## Supplementary material for "Human pluripotent stem cell-derived hepatocyte-like cells for hepatitis D virus studies"

**Supplementary Material Chi et al.**

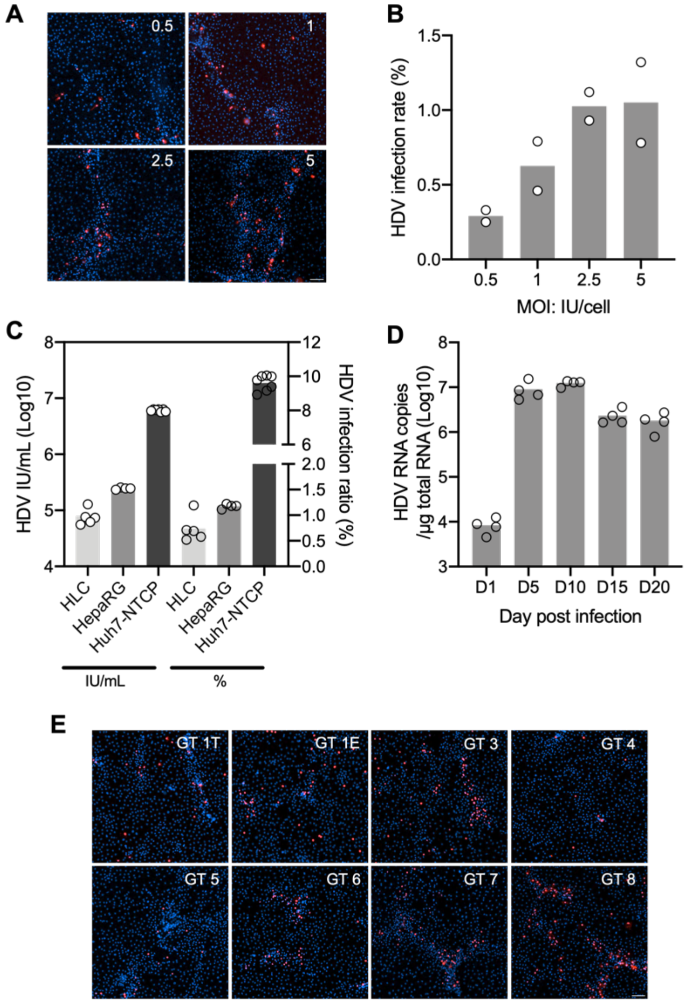


**Figure S1: HLCs are susceptible for HDV infection.** (A & B) HLCs were infected with HDV at different MOIs (0.5, 1, 2.5, or 5) and stained against the HDV antigen (HDAg, red) 5 days p.i. Scale bar = 100 µm. HDAg-positive cells were counted using CellProfiler. (C) HLCs, differentiated HepaRG, and Huh7$^{NTCP}$ were infected with HDV (MOI = 5). (D) HLCs were infected with HDV (MOI = 5) and harvested on indicated days p.i. HDV replication was analyzed by quantifying HDV genome copies in infected HLC lysates using RT-qPCR. Dashed line = LOQ. (E) HLCs were infected with the indicated HDV genotype and 5 days p.i., cells were harvested for HDAg staining. Scale bar = 100 µm.

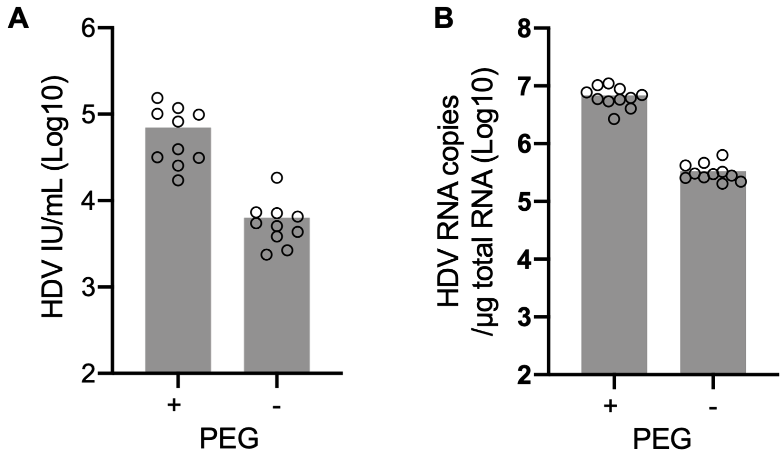


**Figure S2: HDV infection of HLCs is possible in the absence of PEG. (**A & B) HLCs were infected with HDV (MOI=5) in the presence or absence of 4% PEG 8000. Cells were harvested 5 days p.i. and analyzed for HDV infections by counting HDAg-positive cells using CellProfiler (A) or quantifying HDV genome copies by RT-qPCR (B). N = biological replicates.

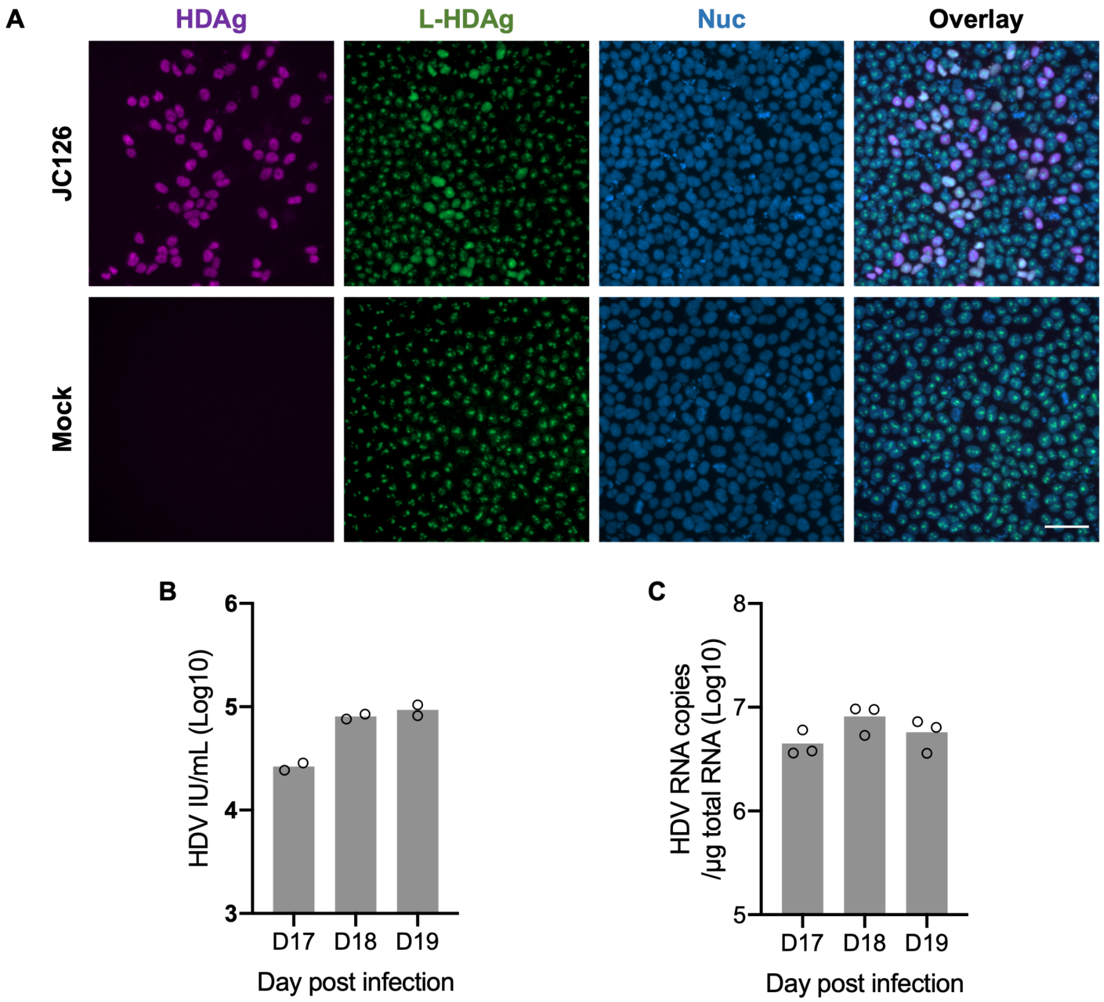


**Figure S3: HDV susceptibility along D17 to D19 of the HLC differentiation protocol.** (A) Stem cells were transfected with or without pJC126. Cells were harvested and assessed by immunofluorescence staining against the HDAg (magenta) and L-HDAg (green) 8 days post transfection. Scale bar = 50 µm. (B-C) HLCs were infected with HDV (MOI = 5) at the indicated day of the differentiation protocol and harvested for analyzing HDV infection efficiency by quantifying HDV-positive cells through CellProfiler (B) or detecting HDV genome copies through RT-qPCR (C) on 5 days p.i.. N = biological replicates.

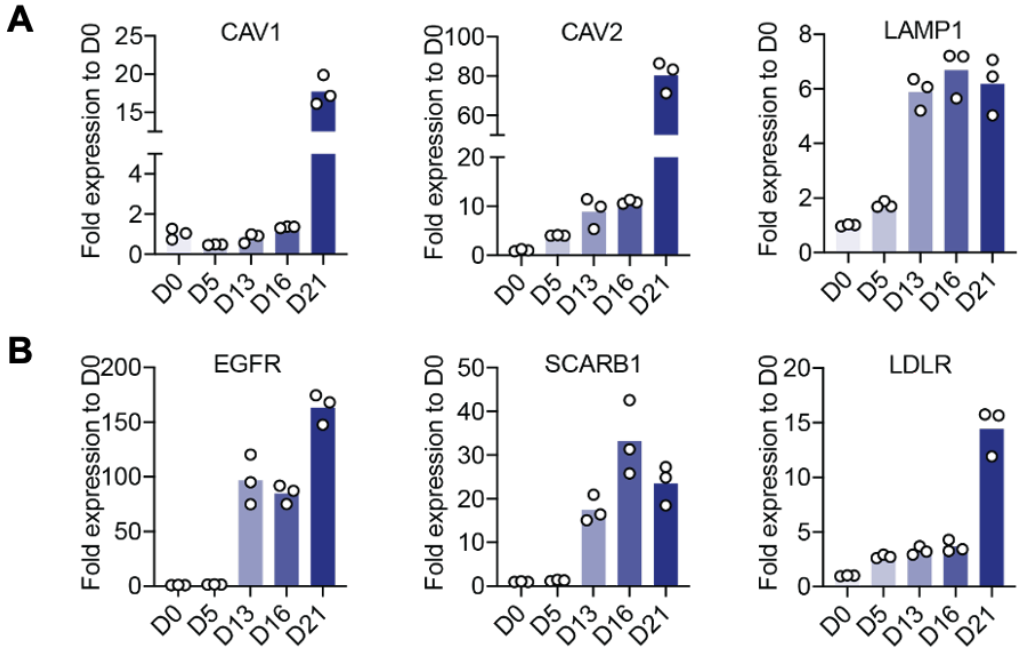


**Figure S4: Gene expression along HLC differentiation.** Total RNA was extracted from HLCs at indicated time points and analyzed for previously described expression of HBV/HDV entry factors (A) and co-receptors (B) expression by RT-qPCR. N = biological replicates.

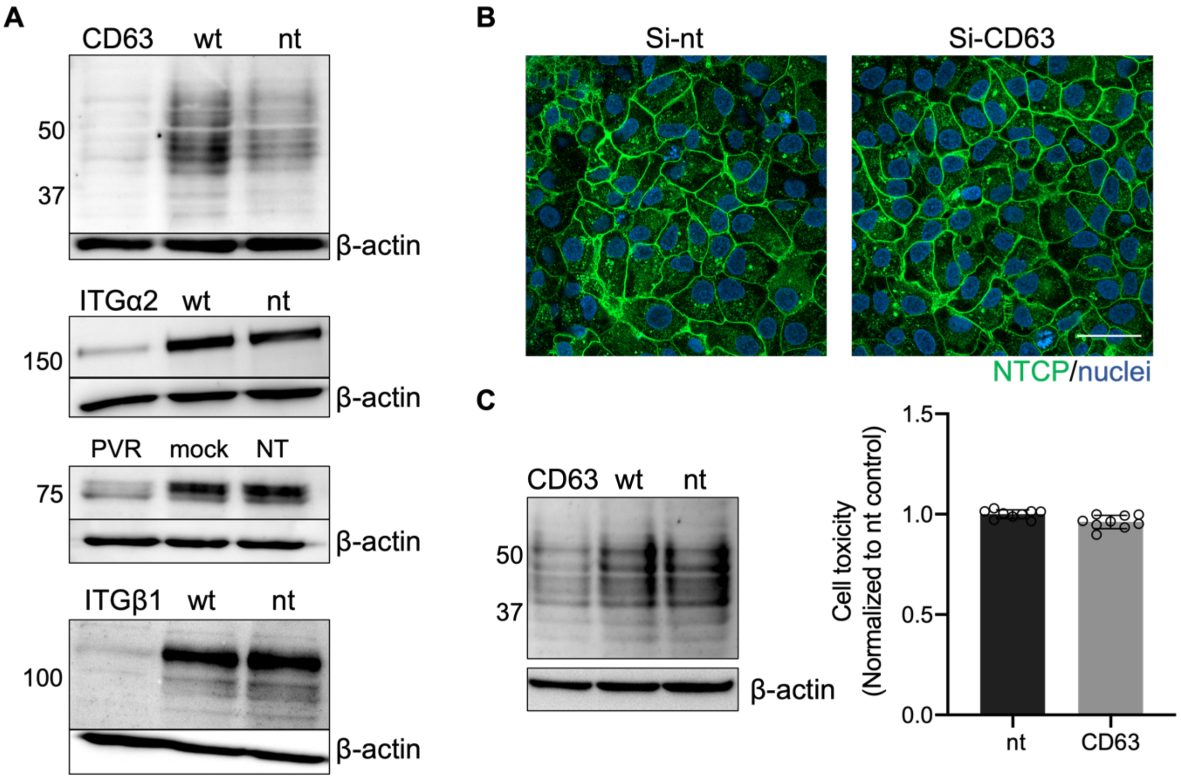


**Figure S5: Knockdown of selected target genes.** Four siRNAs targeting CD63, ITGA2, PVR, and ITGB1 were delivered into Huh7$^{NTCP}$ cells and downregulation was confirmed by western blot analysis on cell lysates harvested 3 days later. Untransfected (wt) and non-target (nt) siRNAs were used as controls. (B) Huh7$^{NTCP}$ were transfected with 50 nM siRNA targeting CD63. 3 days later, cells were stained with Atto-MyrB-488 (green) and imaged by confocal microscopy. Scale bar = 50 µm. (C) HepG2$^{NTCP}$ cells were transfected with 50 nM siRNA targeting CD63. 3 days later, cells were harvested for CD63 expression analysis using western Blot (left) and toxicity assays (right).

### Supplementary Materials & Methods

*Stem cell transfection*

300 ng pJC126 (or Mock) and 3 µl lipofectamine Stem Transfection Reagent (Invitrogen) were each mixed with 25 µl OptiMEM in two tubes. After incubating at RT for 5 min, the content of the two tubes was mixed and incubated for an additional 30 min at RT. WA09 cells were dissociated into single cells using Accutase, and $4x10^4$ cells were seeded in mTeSR1 supplemented with 10 µM Y-27632 on a Matrigel coated well of a 24-well plate. The transfection mix was added dropwise to the cells. Cells were passaged once and then fixed for immunofluorescence analysis nine days after transfection.

*Immunofluorescence (IF) staining and microscopy*

For IF staining, cells were fixed with 4% paraformaldehyde (PFA, Science Services) for 15 min at room temperature (RT). For the staining against HDAg, HBcAg, and HBsAg, cells were permeabilized with 0.5% TritonX-100 (Millipore) for 10 min, followed by the incubation with primary antibody diluted in 1% casein overnight at 4°C. For differentiation marker staining (ALB and AFP) and CD63, cells were blocked and permeabilized in PBTG (10% goat serum, 1% bovine serum, and 0.1% TritonX-100 in PBS) for 30 min prior to incubation with primary antibody diluted in PBTG overnight at 4 °C. After three washes with PBS, cells were incubated with secondary antibodies diluted either in 1% casein or PBTG for 1 h at RT. For NTCP staining, cells were fixed with 1.25% PFA, followed by overnight incubation with 1.5 µM $Atto^{488}$ or $Atto^{565}$ labeled Myrcludex B[75] ($MyrB^{Atto488/565}$) at 4 °C. Hoechst 33342 (1:1000, Thermo Fisher Scientific) was used for nuclei staining. Images were taken either on a Zeiss Cell discoverer 7 (CD7) microscope or a confocal microscope Zeiss Airyscan 2. For the quantification of HDV infection events, roughly 70% of cells grown in a well of a 24-well plate were imaged. Images were processed using ImageJ and ZEN software. HDV infection events were quantified using CellProfiler[76].

*Quantitative reverse-transcriptase PCR (qRT-PCR)*

Total RNA was isolated from cell lysates using the NucleoSpin RNA kit (Macherey-Nagel) and from cell supernatants using the QIAmp viral RNA mini kit (Qiagen) according to the manufacturers' protocols. The secondary structure of HDV RNA was denatured by incubating RNA at 95 °C for 5 min followed by fast cooling to -80 °C. Reverse transcription was performed using the High-Capacity cDNA Reverse Transcription Kit (ABI). HDV RNA was detected using the Luna Universal Probe qPCR Master Mix (New England Biolabs) with the primers and probe listed in **Table S1** on a CFX96 thermocycler (Bio-rad). The JC126 plasmid encoding HDV GT1, kindly provided by Dr. John Taylor, was used to generate a standard for quantifying HDV RNA copy numbers. The expression of other genes was quantified using iTaq™ Universal SYBR ® Green Supermix (Bio-rad) with primers listed in **Table S1**. Both the HDV RNA copies and relative expression data were normalized to expression against the housekeeping gene RPS11.

*Enzyme-linked immunosorbent assay (ELIS*A)

Secreted hepatitis B surface antigen (HBsAg) was quantified in the cell supernatant by enzyme-linked immunosorbent assay (Architect, Abbott) according to the manufacturer's protocol. Absolute values above 0.05 IU/mL were scored positive.

*RNA-seq analysis*

RNA isolated from biological duplicates using the NucleoSpin RNA kit (Macherey-Nagel) was used as input. RNA integrity was determined using Agilent RNA Nano 6000 chips on the Agilent Bioanalyzer 2100 system (Agilent Technologies). Sequencing libraries were constructed using the NEBNext Ultra II Directional RNA Preparation Kit (New England Biolabs) and the NEBNext Multiplex Oligos for Illumina. Samples were sequenced on the Illumina NextSeq 550 (150 cycles) in paired-end mode (paired-end sequencing) at the Deep Sequencing Core Facility of Heidelberg University.

RNA-seq reads were mapped to the human transcriptome assembly GRCh38 release 95 (also known as hg38) using Salmon version 0.15.2[77] to estimate transcript abundance. Estimated abundances were then aggregated at the gene level using tximport version 1.24.0[78]. Data normalization and differential expression analysis was performed using DESeq2 version 1.36.0[79]. Finally, Gene Ontology (GO) enrichment analysis of the set of differentially expressed genes was performed using topGO version 2.48.0[80].

*siRNA reverse transfection*

42 selected SMARTpool siRNAs (Dharmacon) were individually added to each well of a 96-well plate containing 20 µl OptiMEM and 0.2 µL Lipofectamine™ RNAiMAX Transfection Reagent at a final concentration of 12.5 nM for Huh7$^{NTCP}$ and 50 nM for HepG2$^{NTCP}$ cells. Afterwards, 2 x $10^4$ Huh7$^{NTCP}$ cells or 3 x $10^4$ HepG2$^{NTCP}$ cells were added to each well in 100 µl culture medium. 48 h later, the supernatant was replaced with fresh medium and cells were used for downstream experiments 24 h later.

*siRNA forward transfection after HDV infection*

1.5 x$10^4$ Huh7$^{NTCP}$ cells were seeded per well of a 96-well plates in 120 µl culture medium. Cells were infected with HDV (MOI=1) 24 h after cell seeding, followed by siRNA transfection on the next day. 50 nM siRNA and 0.2 µl Lipofectamine™ RNAiMAX were each mixed with 10 µl OptiMEM in 2 tubes. After incubating at room temperature for 5 min, the two tubes were mixed and incubated for an additional 30 min at room temperature. Then, the transfection mix was added dropwise to the cells. After 48 h, the supernatant was removed and replaced with fresh medium. 24h later, the cells were used for downstream experiments, e.g. HDV infections.

*Western blot analysis*

Cells were lysed in RIPA lysis and extraction buffer (Thermo Fisher Scientific) supplemented with a 1X protease inhibitor cocktail (50X cOmpleteTM Mini Protease Inhibitor Cocktail, Roche). Cell lysates were separated by 12% sodium dodecyl sulfate–polyacrylamide gel electrophoresis (SDS-PAGE). Proteins were electro-transferred onto a 0.45 µm polyvinylidene fluoride (PVDF) membrane. The membrane was blocked in PBS containing 5% dry milk (Roth) for 1 h at RT and incubated with primary antibody overnight at 4 ℃. After washing with PBS containing 0.1% Tween 20, the membrane was incubated with horseradish peroxidase-labeled goat-anti-mouse or goat-anti-rabbit antibodies (used at 1:4000, Jackson Immuno Research) for 1 h at RT. The membranes were imaged on the INTASELL Chemostar imager.
